# MHC-II acts as a fusion-triggering receptor for bat influenza virus

**DOI:** 10.64898/2026.07.08.737388

**Authors:** Sarah Peterl, Jonathan Robert, Maria K. Osman, Rebecca A. Haines, Konstantin Fischer, Richard E. Langi, Kevin Ciminski, Martin Schwemmle, Peter Reuther, Petr Chlanda

## Abstract

Influenza A virus hemagglutinin is a prototypical class I viral fusion protein that binds sialylated glycans and is activated by low pH in endosomes. In contrast, bat-derived IAV subtypes H17N10 and H18N11 use major histocompatibility complex class II (MHC-II) as an entry receptor, but how this receptor contributes to membrane fusion remains unknown. We find that MHC-II-dependent hemagglutinin subtypes H17, H18, and H19 possess an increased negative net charge relative to canonical HAs. Using cryo-electron tomography, we demonstrate that H18N11 morphology remains stable and H18 is in prefusion conformation at strongly acidic pH. Remarkably, H18 undergoes fusion-relevant conformational changes only when both MHC-II binding and low pH are present. By reconstitution of H18N11 fusion with liposomes and purified MHC-II, we show that receptor engagement is required to trigger the fusion activity of H18. These findings identify MHC-II as a receptor that directly triggers membrane fusion and reveal a previously unrecognized receptor-dependent mechanism of influenza virus entry.

**Significance:** Bat-derived influenza A viruses (IAV) challenged established paradigms because they use major histocompatibility complex class II (MHC-II) molecules instead of sialylated glycans for cell entry. Recent studies suggest that MHC-II usage extends beyond bats, with avian, swine, and human strains showing dual receptor specificity. However, how MHC-II contributes to membrane fusion has remained unclear. We show that, unlike conventional HAs that are activated by acidity alone, the H18 fusion protein requires receptor engagement and low pH to undergo fusion-relevant conformational changes leading to membrane fusion. Hence, we reveal a novel receptor-dependent mechanism of membrane fusion for the MHC-II dependent entry of H18N11.

## Introduction

Influenza A viruses (IAV) stand out for their broad host range and genetic diversity, presenting a significant threat to global health by breaching the species barrier to humans. Wild aquatic birds are recognized as natural reservoir for most IAV subtypes (1). With the discovery of divergent subtypes H17N10, H18N11, and H18N12 in different New World bat species, the ecological landscape of IAV has been expanded. (2–5). Viral host range is determined by a combination of molecular, cellular, and environmental factors. Essential early steps of IAV infection are receptor binding and membrane fusion. The surface glycoprotein hemagglutinin (HA) subtypes H1 – H16 recognize sialic acid linkages at the target cell surface. It has been shown that major histocompatibility complex II (MHC-II) is an essential entry determinant for New World bat IAV and that H17 and H18 do not bind to sialic acid (6, 7). MHC-II shuttles to lysosome-like compartments, where antigen processing and peptide loading occur before surface presentation. As H17 and H18 can engage a variety of vertebrate MHC-II for entry, including human, there is the possibility of spillover to humans (8), though this risk is considered minor since H18N11 was reported to be poorly adapted to non-bat species (9, 10). In addition to bat IAV, avian H19 with MHC-II specificity, as well as H2N2 and H3N2 viruses with dual MHC-II and sialic acid specificity, have been recently identified (11–13).

In bats, H18N11 was predominantly detected in the follicle-associated epithelium (FAE) of gut-associated lymphoid tissue and, to a lesser extent, in the squamous epithelium of the palatine tonsils, whereas in ferrets, strong viral RNA and antigen signals were observed in the FAE of the pharyngeal and palatine tonsils (10). As these tissues are enriched in antigen-presenting cells, including dendritic cells and macrophages, bat IAVs have adapted to target preferentially MHC-II expressing cells, including professional phagocytes (14).

Early IAV infection steps, including receptor binding, uptake into the cell, pH-dependent activation of HA and fusion with the endosomal membrane are well described for canonical IAV infecting other species than bats (15, 16). HA is a class I fusion protein, which is synthesized as a trimeric protomer (HA0) and subsequently proteolytically cleaved into HA1 and HA2 domains. While the receptor binding site is located in the HA1 domain, the fusion peptide that is inserted into the endosomal membrane is part of the HA2 domain (17). Following receptor binding, the virus is taken up by the cell through endocytosis or macropinocytosis (18). A low pH environment within endosomes is the trigger for major conformational changes in HA. First, a reversible intermediate state of HA is induced. Later, conformation is changed irreversibly, initiating fusion with the endosomal membrane (19, 20). Depending on the viral subtype and host, the fusion pH can vary in a range between 4.8-6.2 within the 16 canonical HA subtypes (21, 22).

H17 and H18 have a monobasic cleavage site. For H17 and H18, which carry engineered polybasic cleavage sites, fusion pH of 5.4 and 5.6 have been reported, respectively (23). In contrast, H17 with its natural monobasic cleavage site, shows increased resistance to low pH (24), suggesting a different entry mechanism of bat IAVs. At the amino acid level, H17 and H18 share an average of 50% identity with conventional IAV HAs and the receptor binding site lacks the typical sialic acid binding cavity and instead harbours acidic residues (25).

While it was shown that conserved amino acids within the a2 domain of MHC-II are required for H18-mediated entry, the MHC-II-binding site of H18 remains elusive (8). A recent study by Osman et al. revealed that binding of H18N11 viral particles to MHC-II on the cell surface induces clustering of MHC-II, possibly to increase the avidity required for attachment. However, structural and mechanistic insights into the subsequent steps of bat IAV entry, including endocytosis and membrane fusion are lacking.

To fill this gap, we have combined cryo-electron tomography and fluorescence spectroscopy. We show that H18N11 is stable at pH 4 and requires receptor engagement to trigger activating conformational changes of HA required for membrane fusion, proposing a new role of MHC-II as a fusion receptor.

## Results

### Bat IAV H18N11 virions incorporate variable proportions of hemagglutinin and neuraminidase

To characterize the morphology of the wild-type H18N11 (H18N11_wt_) virus and to map the distribution of its surface glycoproteins, we performed cryo-electron tomography (cryo-ET) on supernatants harvested from MDCK cells stably expressing human HLA-DR with a titer of 1 × 10^5^ ffu/ml. Surprisingly, H18N11_wt_ infection generated virions with three distinct glycoprotein compositions: particles displaying both HA and NA, particles bearing HA alone, and particles bearing NA alone, either with or without a visible matrix layer (Fig 1B). Strikingly, 33 % of examined particles (n=21) contained NA but lacked any detectable viral ribonucleoprotein complexes (vRNPs) (Fig 1C).

**Figure 1:**
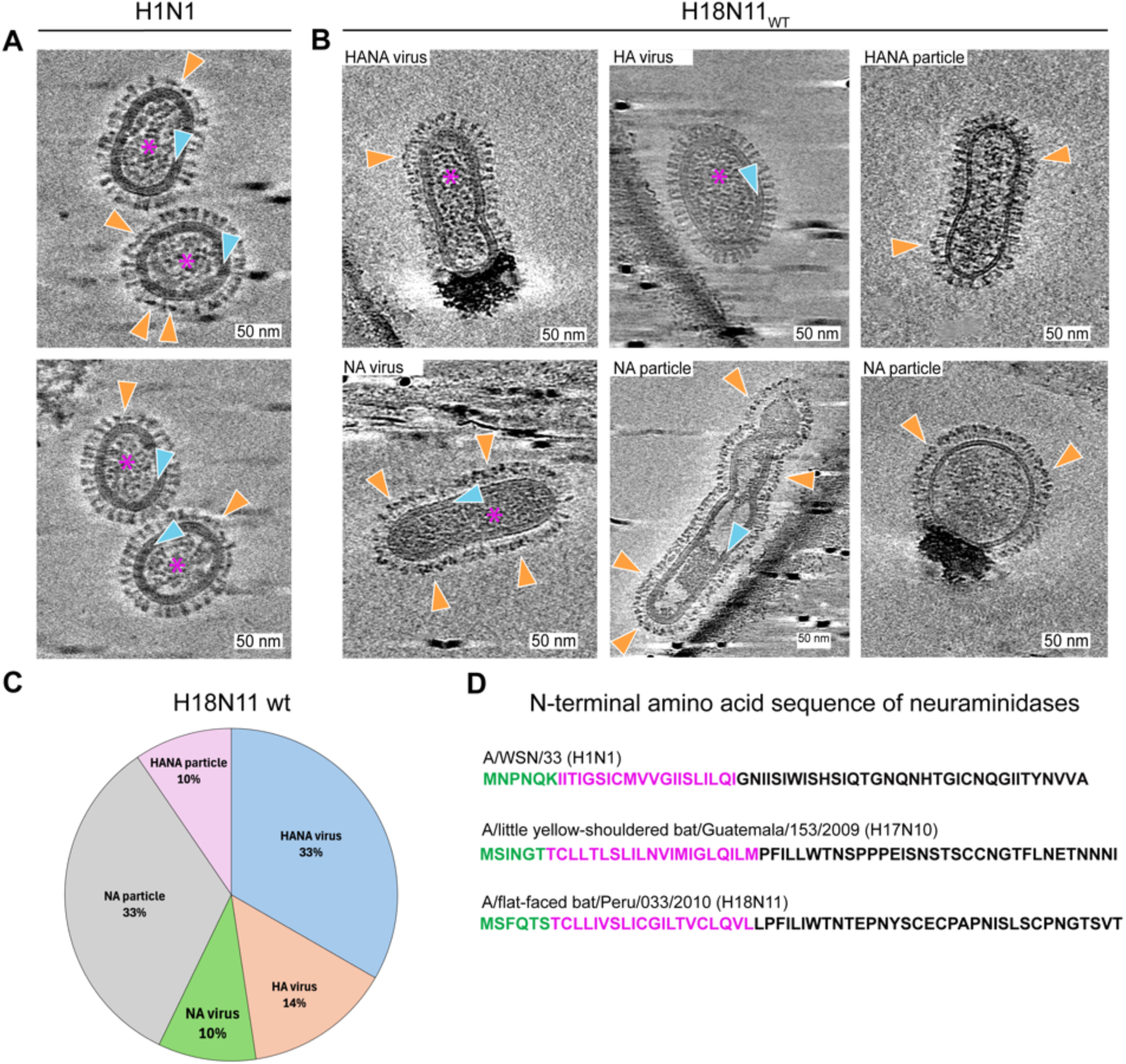
Cryo-electron tomography of H18N11_wt_ viruses. **A.** Slices through a cryo-electron tomogram of A/WSN/33 (H1N1) virions. **B** Slices through a cryo-electron tomogram of particles found in H18N11_wt_ preparations. Exemplary individual NA glycoproteins and NA clusters are indicated by orange arrowheads. M1 layers are indicated by blue arrowheads, vRNPs are indicated by magenta asterisks. **C.** Distribution of different particles (n=21). **D.** Amino acid sequences of the N-terminal region of NA from H1N1, H17N10, and H18N11 viruses. The cytosolic tail and transmembrane domain are highlighted in green and magenta, respectively.

This is in contrast to conventional sialic acid (SA)-dependent IAV, such as the A/WSN/33 (H1N1), which typically harbor both HA and NA on the envelope in an approximate 5:1 HA:NA ratio for spherical particles (26). Moreover, NA in virions is usually clustered at the distal pole of the particle, a distribution that depends on NA cytoplasmic tail (26) and is visible by cryo-ET (Fig 1A). NA can be distinguished from HA by a large head and a lack of visible stem density. Our data show that the N11 head is positioned farther from the viral envelope than N1 in A/WSN/33 (Fig 1A, B, orange arrowheads). This indicates that the N11 stem is longer than N1. Recent work has shown that the NA cytoplasmic tail of sialic acid-dependent IAVs directs NA to the rear of the virion, thereby optimizing its sialidase activity (27). While NA tail (MNPNQK) is highly conserved across conventional IAVs, it differs markedly in the bat-derived N10 and N11 proteins (Fig 1D). Collectively, the heterogeneous HA/NA patterns observed in H18N11_wt_ virions underscore functional distinctions between H18-N11 and the conventional HA-NA pairs of IAVs.

### Bat IAV H18N11 is resistant to low pH

Since IAV cell entry is pH-dependent, we next sought to analyze the pH stability of H18 as well as the overall architecture of H18N11_WT_ virions at acidic pH. The conventional human A/WSN/33(H1N1) served as control. Purified viruses were treated with different pH buffers in the range of pH 4.0-7.5 to mimic endosomal acidification upon cell entry. Viruses were subsequently vitrified by plunge-freezing and analyzed by cryo-ET. Both H1N1 and H18N11_wt_ showed a clearly visible viral envelope with embedded HA trimers in prefusion conformation and an intact matrix layer at pH 7.4 (Fig. 2A, C). While H1N1 virions underwent a conformational switch of HA into postfusion conformation and matrix layer uncoating at pH 5.5, these changes were not observed for H18N11_wt_ (Fig. 2B, D). Remarkably, H18 retained its prefusion conformation even at pH 4. This indicates that low pH alone was not sufficient to induce conformational changes of H18 or viral uncoating of bat IAV H18N11_wt_ necessary for fusion and release of the genome.

**Figure 2:**
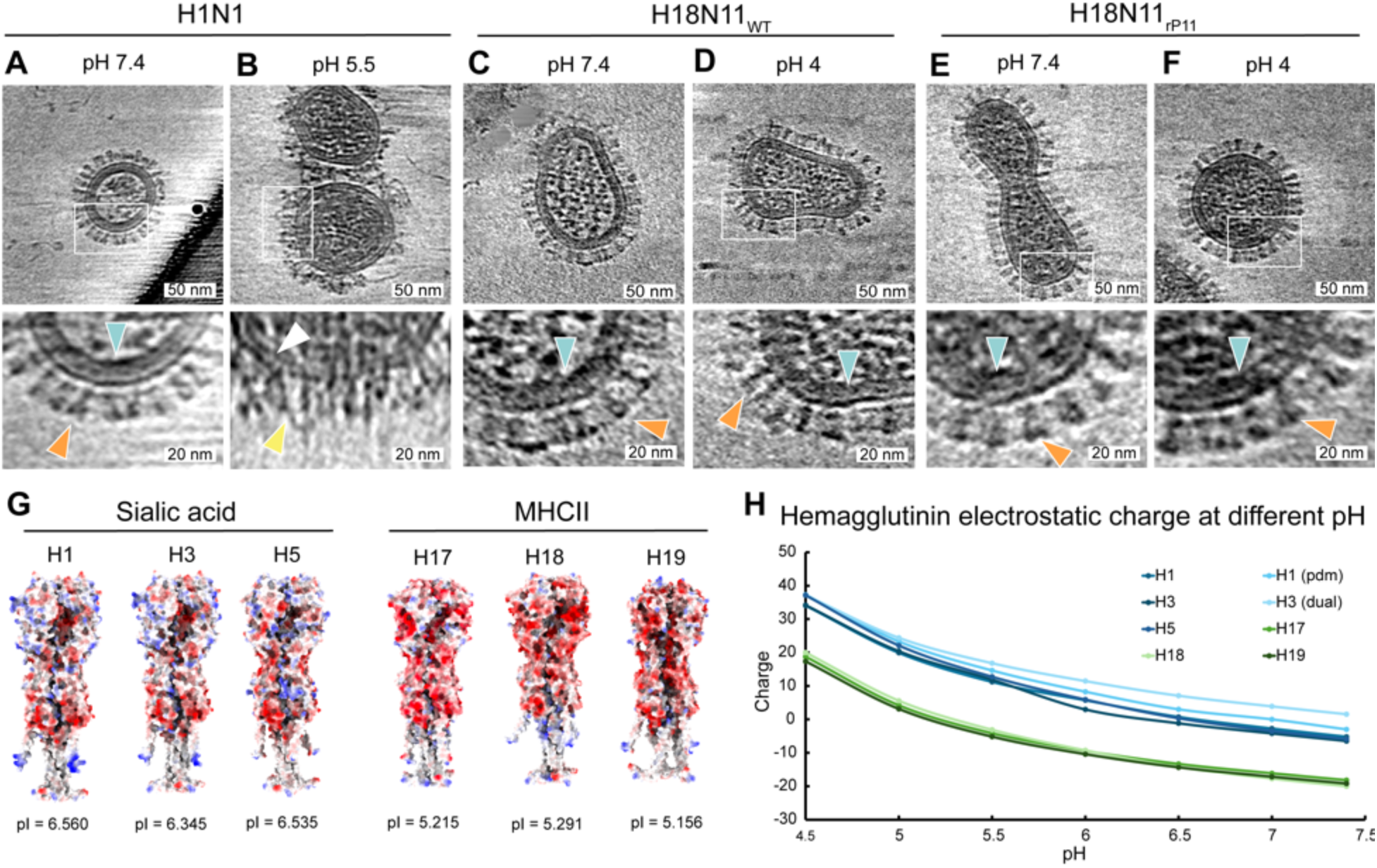
Reduced H18N11 pH sensitivity and electrostatic charge. **A.** Slice through a cryo-electron tomogram of an A/WSN/33 (H1N1) virion at pH 7.4 with a magnified area indicated by a white rectangle showing H1 in prefusion conformation (orange arrowhead) and an intact M1 layer (blue arrowhead) beneath the viral envelope. **B.** Slice through a cryo-electron tomogram of two H1N1 virions at pH 5.5 and zoom-in showing H1 in disorganized postfusion conformation (yellow arrowhead) and the viral envelope without intact M1 layer (white arrowhead). **C.** Slice through a cryo-electron tomogram H18N11_wt_ virion at pH 7.4 with zoom-in showing H18 in prefusion conformation (orange arrowhead) and an intact M1 layer (blue arrowhead) beneath the viral envelope. **D.** Slice through a cryo-electron tomogram of a spherical H18N11_wt_ virion at pH 4 with zoom-in showing H18 in prefusion conformation (orange arrowhead) and an intact M1 layer (blue arrowhead) beneath the viral envelope. **E.** Slice through a cryo-electron tomogram H18N11_rP11_ virion at pH 7.4 with zoom-in showing H18 in prefusion conformation (orange arrowhead) and an intact M1 layer (blue arrowhead) beneath the viral envelope. **F.** Slice through a cryo-electron tomogram of a spherical H18N11_rP11_ virion at pH 4 with zoom-in showing H18 in prefusion conformation (orange arrowhead) and an intact M1 layer (blue arrowhead) beneath the viral envelope. **G.** AlphaFold3 models of exclusive sialic acid- or MHC-II-dependent HAs. **F.** Net charge at different pH for H1 = A/WSN/33 (H1N1), H1(pdm) = A/California/07/2009 (H1N1), H3 = A/Hong Kong/1/68 (H3N2), H5 = A/Texas/37/2024 (H5N1), H3(dual) = A/Victoria/361/2011 (H3N2), H17 = A/little yellow-shouldered bat/Guatemala/153/2009 (H17N10), H18 = A/flat-faced bat/Peru/033/2010(H18N11), and H19 = A/common pochard/Kazakhstan/Kz52/2008. Surface charge displayed in ChimeraX with red showing acid and blue basic residues, respectively. Isoelectric points (pI) and net charge of HAs at different pH were calculated using https://www.protpi.ch/Calculator/ProteinTool.

Since the H18N11_wt_ virus produces low titers, we confirmed the above observations on H18 stability at low pH, using a cell culture-adapted H18N11 variant rP11 (H18N11_rP11_), which produces higher titers (6×10^7^ ffu/ml) in MDCK cells stably expressing human MHC-II HLA-DR. H18N11_rP11_ retains the NA cytosolic tail and transmembrane domain but lost the NA ectodomain via the introduction of a premature stop codon (G107X) in the NA segment and acquired two substitutions in the HA head domain (K170R, N250S) (10). As expected, the NA ectodomain was not observed in any of the analyzed virions (n=152), confirming that NA is truncated and thus absent in H18N11_rP11_ particles (Fig. 2E). Cryo-ET analysis of acidified H18N11_rP11_ (Fig. 2F) confirmed that H18 is stable at low pH as shown for H18N11_wt_.

To explore the pronounced acid-stability of H18, we analyzed the electrostatic surface of an AlphaFold 3 model of H18 and compared it with the surfaces of several other HAs. For each protein, we computed the theoretical isoelectric point (pI) and plotted the net charge as a function of pH. SA-dependent IAV HAs (H1, H3, H5) displayed a markedly different charge distribution than the exclusive MHC-II-dependent H17, H18, and H19, which are overall more acidic (Fig. 2G). This trend was reflected in their pI values: H17, H18, and H19 possess significantly lower pI than the conventional HAs, indicating that a higher proton concentration (i.e., lower pH) is required for these proteins to acquire the same net positive charge (Fig. 2G). Consequently, H18 retains a neutral or slightly negative surface charge under acidic conditions (Fig. 2H), which explains its stability at low pH. The same electrostatic profile was observed for H17 and H19, suggesting that acid resistance is a general feature of exclusive MHC-II-dependent HAs (Fig. 2H).

### MHC-II binds to H18 at low pH and triggers postfusion conformation

As shown by the hemagglutination assay (28), H1N1, but not H18N11_rP11_, agglutinated red blood cells (Fig. 3A, B), consistent with the inability of H18 to bind SA residues (29). To compare membrane fusion efficiency at different pH, we performed cell-cell fusion assays at pH 5.5 and 4.5 by transiently expressing H1 or H18 together with GFP in MDCK-HLA-DRTagRFP cells (Fig. 3C). While H1 induced syncytia formation at pH 5.5, the number of H1-induced syncytia was markedly reduced at pH 4.5, indicating that more acidic pH is suboptimal for H1-mediated fusion (Fig. 3C, D). In contrast, H18 triggered syncytia formation only at pH 4.5 but not at pH 5.5, suggesting that its fusion-relevant conformational changes are adapted to this more acidic environment (Fig. 3C, D). Because low-pH exposure alone failed to drive H18 into its post-fusion conformation, we investigated whether engagement of MHC-II could provide the missing trigger for HA rearrangement. We used the cell culture-adapted H18N11_rP11_ virus, which displayed the same acid-stability as H18N11_wt_. H18N11_rP11_ particles were mixed with soluble HLA-DR ectodomains and subjected to pH 7.4 or pH 4.5. The complexes were then plunge-frozen and examined by cryo-electron tomography to assess HA conformational states under each condition. A soluble non-classical MHC-II molecule HLA-DM, which does not support H18N11 entry, served as a control (8). At neutral pH, few H18:HLA-DR binding events were observed (Fig. 3E), indicating a weak binding affinity between the two proteins. Remarkably, when H18N11_rP11_ virions were incubated with HLA-DR and subjected to late endosomal pH of 4.5 for 30 min, HA underwent major rearrangements, indicating a switch to postfusion conformation (Fig. 3F, Fig S2). Postfusion HA conformation was characterized by a thin stem domain and a dissociated flexible head domain, composed of HA1 and presumably also HLA-DR ectodomain. The width measurement of the stem domain diameter 2.1±0.3 nm (n=10) was consistent with the width (2-2.5 nm) of the HA2 stem in postfusion conformation (30) and was thinner than the width of HA in prefusion conformation (4.8 ± 0.5 nm, n=10) when measured in tomograms (Fig. 3E, F). Clustering of postfusion HA molecules and areas that were devoid of viral glycoproteins were observed (Fig. 3F). These clusters were often observed in regions of increased viral membrane curvature (Fig. S2). In contrast, no binding or conformational change of HA was observed in the presence of non-canonical HLA-DM (Fig. 3G, H). In conclusion, a combination of acidic pH (4.5) and HLA-DR is necessary to trigger the fusion-competent conformation of H18.

**Figure 3:**
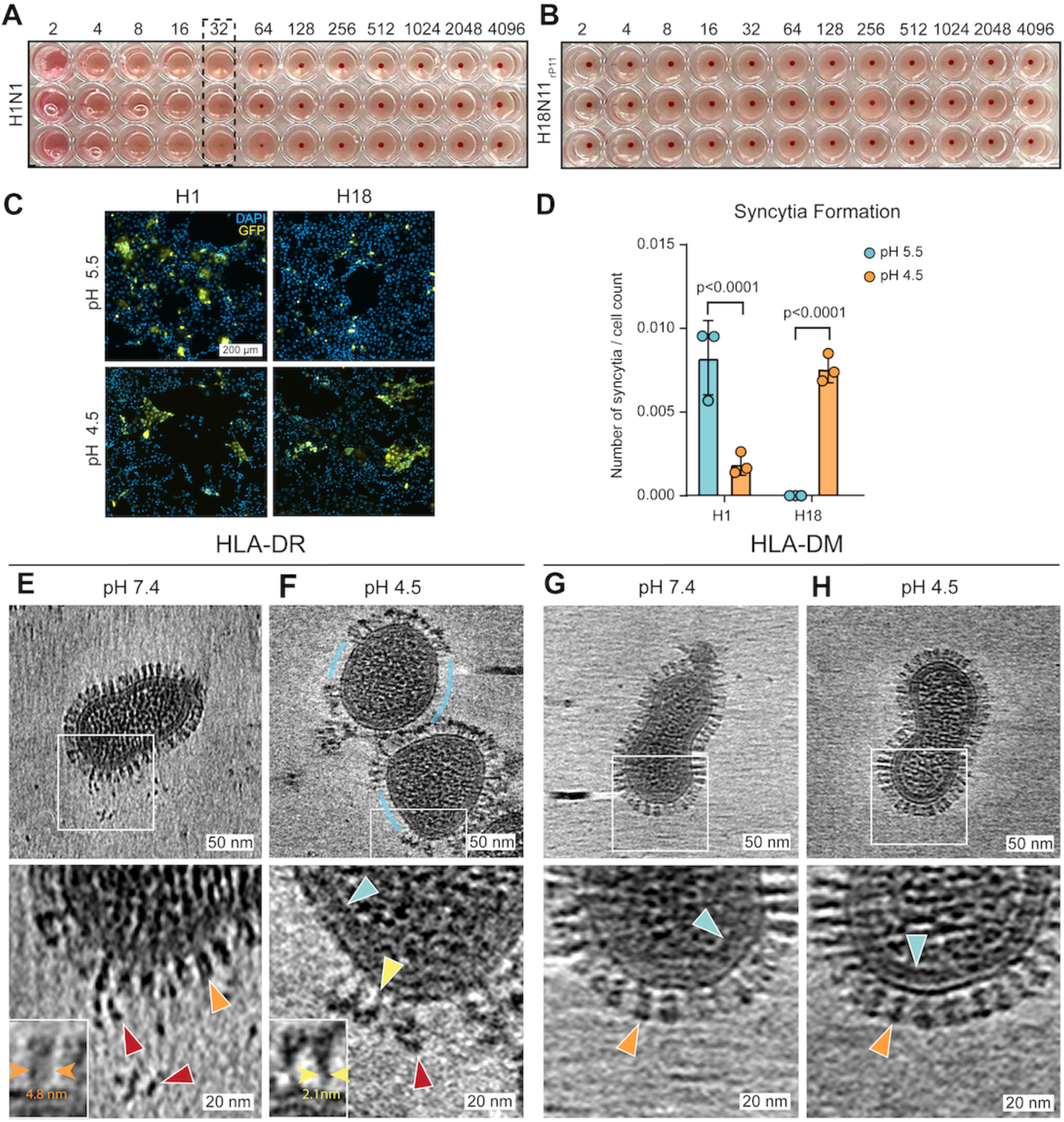
Cryo-ET of H18N11_rP11_-MHC-II interaction and conformational switch. **A.** HA assay of serial two-fold dilutions of A/WSN/33 (1×10^7^ PFU/ml) in three technical replicates showing a titer of 32, indicated by a dotted line. **B.** HA assay of serial two-fold dilutions of H18N11_rP11_ (1×10^7^ ffu/ml) in three technical replicates showing no agglutination. **C.** Representative fluorescence microscopy images of syncytia formation assays on MDCK-HLA-DR_TagRFP_ cells. Cells were co-transfected with expression plasmids encoding GFP and H1 or H18. At 24 h posttransfection, cells were treated with TPCK-trypsin to cleave HA and fusion was induced by the addition of medium adjusted to pH 5.5 or 4.5 for 20 min. The medium was subsequently replaced with neutral pH medium and cells were fixed with 4% PFA after 1.5 h. Nuclei were stained with DAPI. Scale bar for all images 200 µm. **D.** Quantification of syncytia per total cell number per fluorescence microscopy image for the indicated HAs at pH 5.5 (blue) or pH 4.5 (orange). Data points and standard deviations from three independent experiments are shown. Statistical significance was determined by unpaired t-test. **E.** Slice through a cryo-electron tomogram of an H18N11_rP11_ virion incubated with HLA-DR at pH 7.4 with zoom-in showing HA in prefusion conformation (orange arrowhead), soluble and bound HLA-DR (red arrowheads). The inset shows an exemplary HA in prefusion conformation, with a diameter marked by two arrowheads. **F.** Slice through a cryo-electron tomogram of two H18N11_rP11_ virions incubated with HLA-DR at pH 4.5 with zoom-in showing an intact matrix layer (blue arrowhead), H18 in postfusion conformation (yellow arrowhead) and bound HLA-DR (red arrowhead). Areas without glycoproteins between clusters of glycoproteins are marked by blue arcs. The inset shows an exemplary HA in postfusion conformation, with a diameter marked by two arrowheads. **G.** Slice through a cryo-electron tomogram of an H18N11_rP11_ virion incubated with HLA-DM at pH 7.4 with zoom-in showing an intact matrix layer (blue arrowhead) and HA in prefusion conformation (orange arrowhead). **H.** Slice through a cryo-electron tomogram of an H18N11_rP11_ virion incubated with HLA-DM at pH 4.5 with zoom-in showing an intact matrix layer (blue arrowhead) and H18 in prefusion conformation (orange arrowhead). Scale bars: 50 nm, scale bars of zoom-ins: 20 nm.

### MHC-II and low pH are required to trigger H18N11 membrane fusion

Building on these findings, we next investigated H18N11 membrane fusion using liposomes decorated with HLA-DR (Fig. 4A, Fig. S4). Liposomes were composed of 52.5% 1,2-dioleoyl-sn-glycero-3-phosphocholine (DOPC), 40% cholesterol and 5% functionalized DGS-Ni-NTA lipids, which enabled coupling of HLA-DR via a His-tag (Fig S4). In addition, the lipophilic dye 1,1′-Dioctadecyl-3,3,3′,3′-tetramethylindodicarbocyanine, perchlorate (DiD) was incorporated at self-quenching concentration (2.5%) for subsequent analysis by fluorescence spectroscopy (Fig. 4B). While this DiD dequenching does not allow for measuring full membrane fusion, it allows for detecting lipid mixing, which occurs during the hemifusion intermediate (31). Cryo-ET analysis of liposomes confirmed production of unilamellar liposomes in a homogeneous size range and successful binding of HLA-DR (Fig. S4). To monitor H18-liposome membrane fusion over time, we performed fluorescence spectroscopy of H18N11_rP11_ and liposomes with or without HLA-DR at pH 4.5, 5.5, and 7.4 and measured fluorescence dequenching of the lipophilic DiD dye (Fig. 4C). Our results showed that virus-liposome membrane fusion only occurred at pH 4.5 and in the presence of HLA-DR but not at pH 5.5 or 7.4 or in the absence of HLA-DR (Fig. 4C). Taken together, MHC-II is not only required for H18 binding and conformational switch but also membrane fusion in a pH dependent manner. To directly visualize individual steps of viral fusion and confirm that full membrane fusion can take place, we next performed cryo-ET of H18N11_rP11_ incubated with HLA-DR-decorated liposomes under neutral (7.4) or low (4.5) pH conditions (Fig. 4A-E). The fusion process was classified into four distinct phases described previously (32): (I) binding of H18 to HLA-DR, (II) liposomal membrane pinching, (III) formation of an extended interface between H18N11_rP11_ and liposome membranes, (IV) postfusion (Fig. 4A-E). Quantification of contact events at pH 7.5 and 4.5 showed that direct membrane-membrane contact events (class II and III events) were more abundant at low pH (together >70%) compared to neutral pH (20%) (Fig. 4F). While liposomal membrane pinching was also observed at neutral pH, postfusion products were only observed at pH 4.5, not at pH 7.4. In addition, binding events were reduced from approximately 80% at neutral pH to about 20% at low pH (Fig. 4F). Consistent with results from DiD fluorescence dequenching, these data further demonstrated that H18 requires MHC-II and low pH to perform both hemifusion and fusion pore formation.

**Figure 4:**
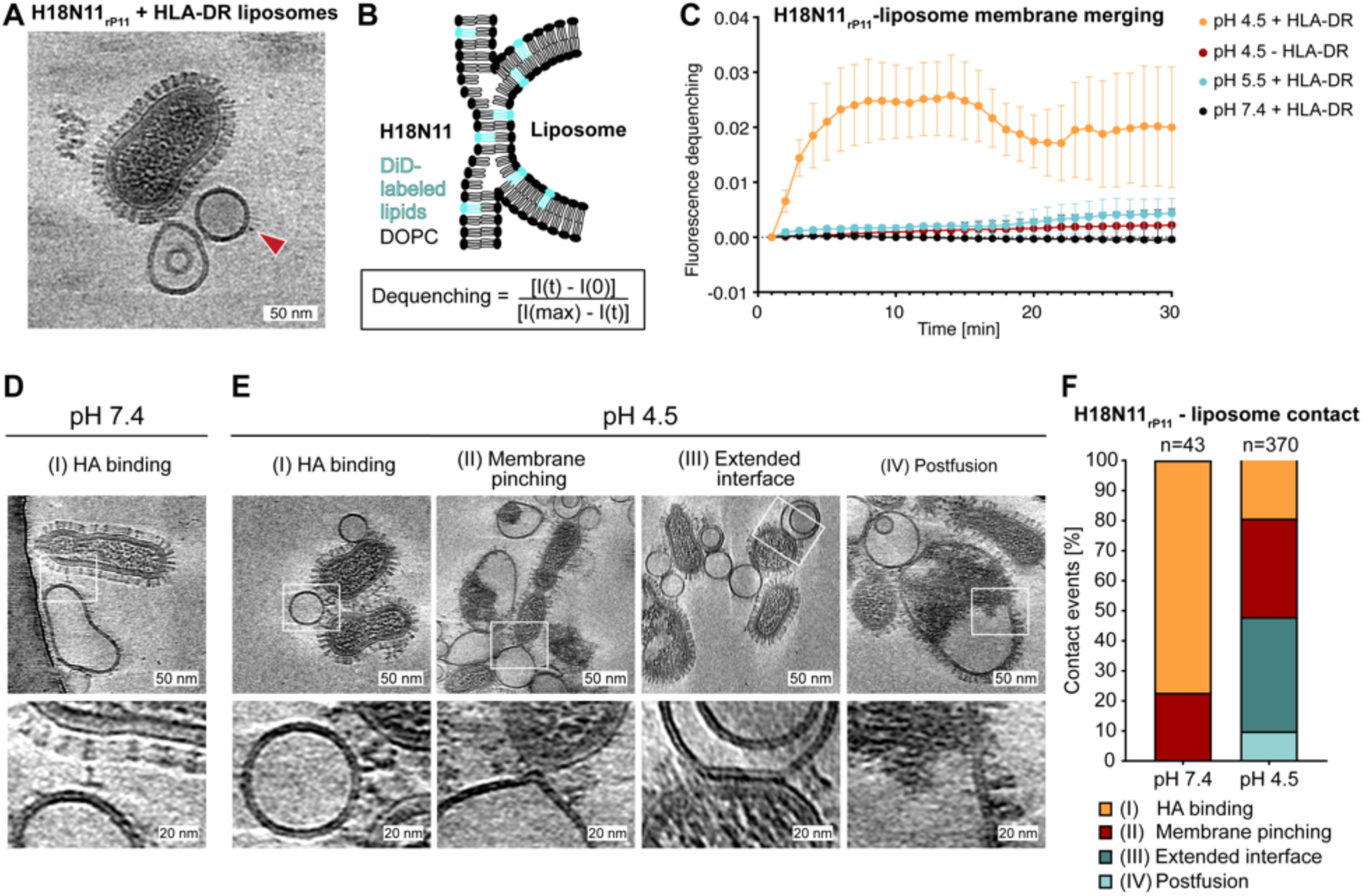
Kinetics and cryo-electron tomography of membrane fusion of H18N11_rP11_ with liposomes functionalized with HLA-DR. **A.** Slice through a cryo-electron tomogram of an H18N1_rP11_ virion and two liposomes decorated with HLA-DR (red arrow) at pH 7.5. Scale bar 50 nm. **B.** Schematic representation of fluorescence dequenching upon virus-liposome membrane merging (hemifusion). Dequenching = (I(t)-I(0))/(I(max)-I(t)) with I(t) = intensity at a given time, I(0) = intensity at t = 0, I(max) = maximum intensity after addition of 5.5 % Triton-X100. **C.** Fluorescence spectroscopy analysis of DiD dequenching during H18N11_rP11_-liposome membrane merging at indicated pH conditions in the presence or absence of HLA-DR relative to the maximum dequenching intensity. Measurements were performed at 37°C every 60 s over a time span of 30 min for three biological replicates. Mean values and standard deviations are indicated. **D.** Cryo-ET slice of an H18N11_rP11_ virion and an HLA-DR coated liposome at pH 7.4 with zoom-in showing H18-HLA-DR binding. **E.** Series of cryo-ET slices showing interaction events between H18N1_rP11_ virions and an HLA-DR coated liposome at pH 4.5 and corresponding zoom-ins. **F.** Quantification of H18N11-liposome contact events at pH 7.5 and 4.5 classified into HA binding (orange), membrane pinching (red), extended membrane interface (dark blue), and postfusion (light blue).

## Discussion

The bat IAVs subtypes H17N10, H18N11, H18N12, and H19 engage MHC-II as entry receptors and encode NAs that lack sialidase activity, representing a fundamental difference from the HA-NA biology of conventional IAVs. For cellular entry, H18N11 can use MHC-II orthologs from various species, including human leukocyte antigen complex HLA-DR (6). We demonstrate that, unlike the conventional A/WSN/33 (H1N1), bat H18N11 virions contain variable amounts of HA and NA (Fig. 1A, B). Notably, N11 lacks the conserved cytosolic tail present in canonical NA (Fig. 1D), which has been recently shown to dictate the localization and HA:NA ratio in canonical IAV (27). Hence, NA in H18N11 is not targeted to the rear end of the virions as in the canonical IAVs. This is further supported by our data showing that approximately one-third of the released particles contain NA and, in addition, lack viral ribonucleoproteins (Fig. 1C). The biological significance of NA-containing particles remains to be elucidated.

IAV, like other enveloped viruses, relies on fusion with target cell membranes to release their genome into the cytoplasm for replication. While some viruses, such as paramyxoviruses (measles and Nipah virus), herpesviruses or retroviruses (HIV-1), directly fuse at the host cell plasma membrane (33, 34), others, e.g. orthomyxoviruses, flaviviruses and filoviruses rely on a low pH environment inside early or late endosomes to fuse with endosomal membranes (35–37). During endosomal maturation, the pH decreases from 6.5 in early endosomes to 5.5 in late endosomes to below 4.5 in lysosomes (Fig. 5) (38, 39).

**Figure 5.**
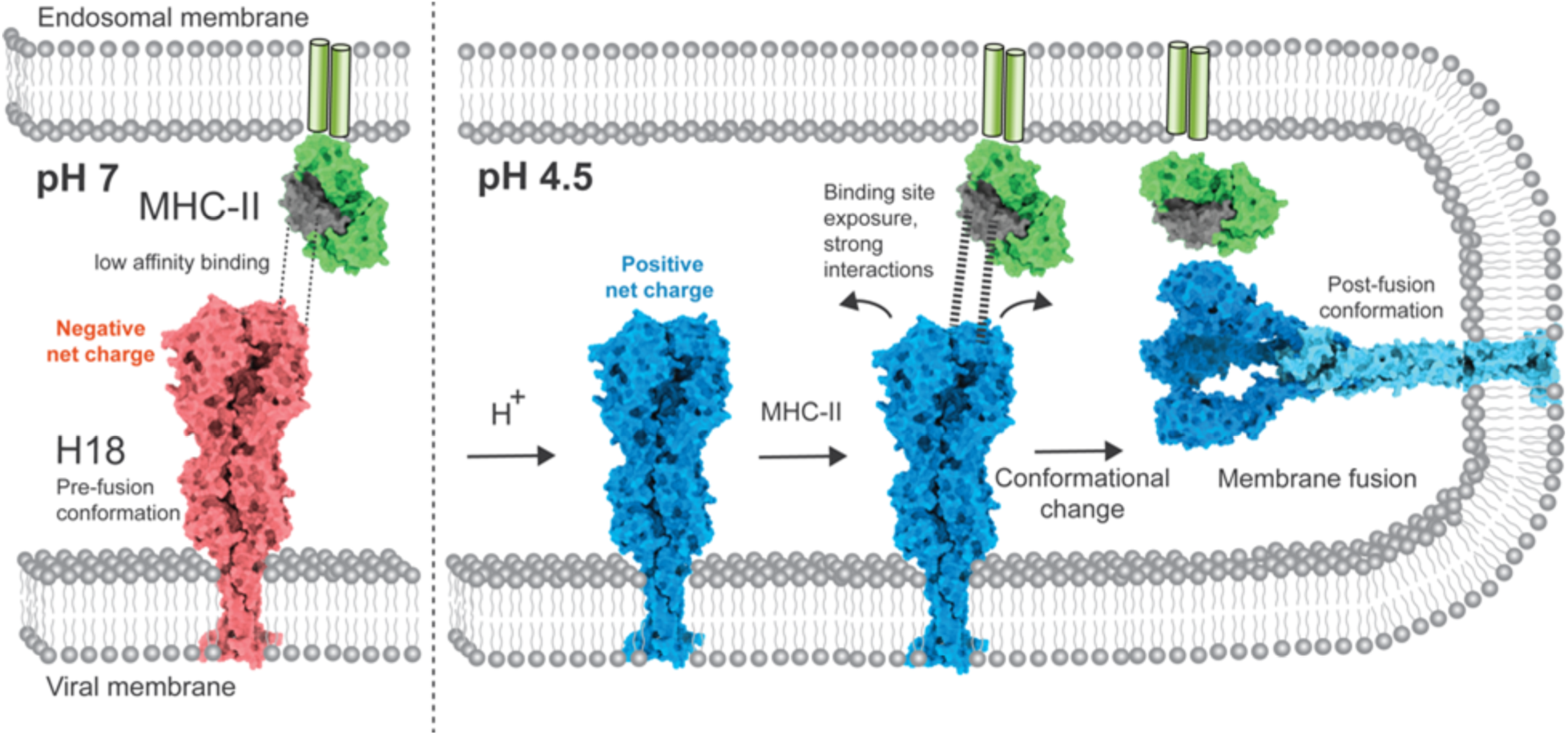
Model for the concerted action of low pH and MHC-II in triggering conformational changes of H18 leading to membrane fusion. H18 (pink) attaches to the target cell surface through low-affinity binding with the alpha 2 domain (dark grey) of MHC-II (green) at neutral pH, which may be enhanced by receptor clustering (47). Exposure to low pH (H^+^) during endosomal or phagosome maturation alone is insufficient to trigger H18 conformational changes (blue). Instead, acidic pH increases the net charge and presumably also small conformational changes, such as dilation of the HA1 head domain (48), and further exposes the MHC-II interaction site. Engagement of MHC-II under these conditions overcomes the energetic barrier required for the transition of H18 into its post-fusion conformation, thereby triggering membrane fusion. HA2 domain is shown in light blue. Used models: Prefusion H18 (AlphaFold3 prediction), MHC-II (PDB: 1AQD, (53)). H18 post-fusion model (hybrid model using post-fusion HA2: PDB:1QU1,(54)).

The pH threshold for IAV fusion ranges between pH 6.2 and 4.8 (21, 22). While avian IAV subtypes are typically activated in a higher pH range (>5.5) (40), a low fusion pH (5.5-4.8) is associated with human adaptation (41, 42). Here, we investigated the morphology and sensitivity of bat IAV H18N11_wt_ and H18N1_rP11_ at pH 7.4, 5 and 4 using cryo-ET and show that H18N11 viruses exhibited remarkable acid stability at pH 4. This finding is in accordance with previous studies reporting resistance of H17 HA at low pH (24). The difference in fusion pH described by Moreira et al., (pH 5.6) may be related to the artificially introduced polybasic cleavage site, which could be linked to fusion activation at higher pH. The conformational switch of HA is a highly regulated process that depends on protonation of amino acids that can act as so-called “pH sensors” including histidines, aspartates, and glutamates (43). Our *in silico* data show that the exclusive MHC-II-dependent H17, H18, and H19 proteins are more acidic (redistributed acidic electrostatics) than their sialic acid-dependent counterparts. A high negative net charge will likely contribute to stability at low pH by stabilizing metastable prefusion conformation via favorable salt bridges and stabilizing interdomain interactions. Previous studies identified histidines at positions 38 and 184 (H3 numbering) as important pH-sensors, which are highly conserved in most IAV subtypes (44, 45). These residues are not present in bat IAVs (Fig. S1) and could additionally contribute to the pH stability of H18N11. Additional to H18 pH stability, we report overall increased H18N11_wt_ and H18N11_rP11_ virion particle stability at low pH (Fig. 2D, F). This may be attributed to the unusual structural and functional characteristics of the M2 proteins of bat IAVs. Thompson et al. revealed reduced ion channel activity of bat M2, which likely contributes to delayed viral uncoating (46).

Given the low receptor affinity, local clustering of MHC-II beneath the virus particle was proposed to be essential for virus attachment and entry (47). Our cryo-ET data on H18N11_rP11_ incubated with an excess of HLA-DR at neutral pH show sporadic binding, supporting recent findings on the low affinity H18:HLA-DR interaction (Fig. 3E). However, when exposed to low pH, HLA-DR induces a conformational switch of H18 from pre- to postfusion (Fig. 3F), leading to membrane fusion (Fig. 4). Based on these findings, we propose that MHC-II acts as a low-affinity receptor and its additional function is to serve as a fusion-triggering receptor for H18. While the MHC-II binding site has been mapped to the *_α_*2 domain of MHC-II dimer (8), the exact binding site at H18 remains to be elucidated. However, we suggest that H18 and MHC-II interactions in the endosome could be enhanced by neutralization of negative charge by protonation (positive net charge), as shown in Fig. 2H. In addition, previous cryo-EM studies have shown that HA trimer can reversibly open (HA1 domain dilation) at low pH intermediate low pH conformation (20, 48), which may expose a structurally optimized MHC-II binding site in the H18 trimer interface.

Interestingly, H3 strain A/Victoria/361/2011 (H3N2), which displays dual-receptor specificity (49), exhibits an isoelectric point (pI) comparable to that of canonical influenza A viruses (Fig. 2H). Consequently, H3 likely retained the pH-sensitivity characteristic of conventional IAVs despite its ability to engage MHC-II as an alternative receptor and thus differs from H17, H18, and H19 lacking sialic acid binding activities. However, it remains to be shown whether the interaction site differs between HAs with dual receptor specificity and HAs with MHC-II specificity.

Taken together, our data suggest that H18N11 virions do not show typical HA-NA distribution as in conventional IAV. Importantly, H18N11 uses a distinct entry mechanism in which MHC-II also acts as a fusion receptor (Fig 5). The findings presented in this study are in contrast to the commonly accepted perception that IAV HA conformational change is triggered solely by low pH (21). Rather, the mechanism of H18N11 entry into host cells is comparable to avian leukosis virus and Ebola virus, which also require receptor interaction to trigger their glycoproteins for subsequent pH-triggered fusion (50, 51) or Lassa virus that relies on a pH-dependent receptor switch (52). We postulate that, as a result of its immune cell tropism (14), H18N11 has adapted to exploit cellular uptake pathways that are typically associated with antigen sampling and degradation in acidic phagolysosomes. Rather than relying solely on conventional receptor-mediated entry at the cell surface, H18 engages MHC-II expressing antigen-presenting cells and remains stable within highly acidic endolysosomal compartments. In this environment, MHC-II interaction appears to act as a trigger for fusion, enabling the virus to escape a pathway that would otherwise lead to degradation and antigen presentation. This strategy suggests an evolutionary adaptation of H18N11 to specifically target professional phagocytes and to repurpose immune surveillance pathways as a niche for productive entry.

## Materials and Methods

### Virus production

The recombinant H18N11_wt_ and the cell culture-adapted H18N11_rP11_ were generated in HEK293T cells using the pHW2000 rescue system (10). HEK293T cells were transfected with eight pHW2000 plasmids comprising the sequence of the respective A/flat-faced bat/Peru/033/2010 (H18N11) segment and four pCAGGS plasmids encoding the PB2, PB1, PA, and NP of the same virus. Cells were transfected with Lipofectamine 2000 following the manufacturer’s instructions. Rescued viruses were propagated for 72 h in MDCKII cells stably expressing HLA-DR (8) and cell supernatants were collected and ultracentrifuged at 77000 g for 2 h on a 30 % sucrose cushion. Finally, viruses were resuspended in DMEM supplemented with 0.2 % BSA, and 1 µg/ml TPCK-trypsin, and stored at -80 °C.

Viral titers were determined via immunofocus assay. MDCKII cells expressing MHC-II were infected and covered with 500 μl of overlay (DMEM, 20 mM HEPES pH 7.5, 200 μg/ml bovine serum albumin (BSA), 20 μg/ml DEAE-Dextran, 100 μg/ml NaHCO_3_, 40 μg/ml oxoid agar) and incubated at 37 °C for 48 h. After overlay removal, cells were fixed with 4 % PFA for 20 min at room temperature, permeabilized with PBS containing 0.1 % Triton, and blocked with PBS containing 5 % FCS. Cells were then stained with a monoclonal anti-H18N11-NP antibody (mouse, produced in-house, 1:500) for 1 h at room temperature, washed and incubated with a secondary HRP-conjugated anti-mouse antibody (Dianova, 1:500) for 1 h at room temperature. After washing, focus-forming units were visualized by adding a substrate solution (PBS supplemented with 0.5 μg/ml 3,3’-diaminobenzidine, 0.5 μg/ml nickel ammonium sulfate and 0.015 % H_2_O_2_). Titers obtained were 1.10^5^ focus forming units (ffu)/ml for H18N11_WT_ and 6.10^7^ ffu/ml for H18N11_rP11_.

### HLA-DR and HLA-DM production

To generate soluble HLA-DR and HLA-DM coding plasmids, sequences for both the α and β chains were synthesized and cloned into the pCDNA3.1 vector using NotI and EcoRI restriction sites. The α-chain constructs include a signal peptide (MDWTWRVFCLLAVAPGAHS), HLA-DRA*0101_1-181_ or HLA-DMA_1-183_, a FLAG tag, a Fos zipper domain, and a 10xHis-tag. The β chain construct comprise a signal peptide (MVLQTQVFISLLLWISGAYG), a Strep-II tag, the CLIP peptide, HLA-DRB1*0101_1-190_ or HLA-DMB_1-200_, a FLAG tag, and a Jun zipper domain. Sequences were synthesized by Azenta Life Science.

Soluble HLA-DR and HLA-DM were expressed in Expi293F cells (ThermoFisher) by transfecting 50 µg of pCDNA3.1_HLA-DRA and 50 µg pCDNA-HLA-DRB1 or pCDNA3.1_HLA-DMA and pCDNA3.1_HLA-DMB for 100 mL of cells at 2.10^6^ cells/ml. Transfection was performed using the ExpiFectamine 293 Transfection Kit (ThermoFisher) following the manufacturer’s instructions. 6 days post-transfection, cells were centrifuged for 20 min at 6000 rpm and the supernatant was filtered with a 0.45 µm filter (Sartorius). Proteins were purified via batch affinity chromatography using HisPur Cobalt Resin (ThermoFisher) and Strep-TactinXT 4 Flow High-Capacity resin (IBA Lifesciences), followed by size exclusion chromatography on a Superdex 200 Increase column into a 20 mM Tris, 150 mM NaCl, pH 8 buffer.

### Liposome preparation

To reconstitute the fusion between bat IAV H18N11 and endosomes in vitro, fluorescently labelled liposomes carrying HLA-DR were produced. Lipids were dissolved in chloroform to produce 100 mM stocks. Lipids were mixed according to Table S1. Chloroform was evaporated under a chemical fume hood using an argon flow for 5 min. The lipid mixture was placed in a vacuum desiccator for 1h. Meanwhile, the reconstitution buffer (20mM Tris, 150 mM NaCl, 10% glycerol) was prewarmed to 41°C. Lipids were dissolved and vortexed for 5 min before performing 12 freeze-thaw cycles in liquid nitrogen and 40-50°C warm water. Extrusion of large unilamellar vesicles was performed using the Avanti Mini-Extruder with a pore size of 100 nm. The extruder was assembled according to instructions provided by the manufacturer and placed on a heating block for 15 min at 42°C. The sample was loaded into the syringe and passed 11 times through the membrane to end in the alternate syringe. Liposomes were stored at 4°C.

Purified HLA-DR (3 mg/ml) proteins were bound to liposomes by incubation in a protein-liposome ratio of 1:100 for 1h at RT on a shaker at 300 rpm to allow interactions between the His-tag and DGS-NTA(Ni) lipids (Avanti Research, 790404P). Unbound proteins were removed in three washing steps with HisPur^TM^ Ni-NTA magnetic beads (Thermo Scientific, 88831). The protein-liposome mix was incubated with beads (1:2 ratio) for 15 min at RT on a shaker (300 rpm). Then, beads with bound proteins were captured on a magnetic stand.

### Fluorescence spectroscopy

Membrane merging between liposomes and H18N11 virions and thus dequenching of the DiD lipid dye was analyzed by fluorescence spectroscopy using a Jasco FP-8500. Virus and DiD-labeled liposomes were incubated in a 1:10 ratio in 100 µl cuvettes. To initiate a fusion reaction, the pH was adjusted to 7.5, 5.5 or 4.5 using liposome buffer (20 mM Tris, 150 mM NaCl, 10% glycerol) adjusted to pH 3 using citric acid. Fluorescence spectroscopy was carried out over a time course of 30 min at 37°C and fluorescence reading at l_Ex_=644 nm, l_Em_= 665 nm every 60 seconds. After 30 min, Triton X-100 (final concentration 5%) was added, and fluorescence reading was continued for 5 min every second. Dequenching was calculated as follows: [F(t) – F(0)]/[F(max) – F(0)], with F(0) being the fluorescence reading before acidification, F(t) the fluorescence reading at a given time point and F(max) the fluorescence reading after Triton addition.

### Cryo-electron tomography

To structurally characterize H18N11 pH stability, H18N11-MHC-II binding and fusion with liposomes, we performed cryo-ET. Viruses were incubated either without MHC-II, with HLA-DR, HLA-DM or liposomes at pH 7.4 or 4.5 for 30 min at RT. A suspension of Protein A-coated colloidal gold (10 nm diameter) was mixed with the samples in a 1:10 ratio shortly prior to plunge freezing. For all conditions, 3.5 µl sample was applied onto a plasma-cleaned EM grid (Qunantifoil R2/1, holey carbon film, Cu, 200 mesh). Plunge-freezing was carried out using an automated Leica EM GP2 plunge freezing device under the following conditions: cryogen: liquid ethane, chamber temperature: 25°C, humidity: 80%, back-side blotting: 1-2 s. Cryo-ET was performed on a Titan Krios Transmission Electron Microscope (TEM, ThermoFisher Scientific) operated at 300 keV, equipped with a BioQuantum LS energy filter with a slit width set to 15eV and a K3 direct electron detector (Gatan), using SerialEM (55). Overview maps at 8700x magnification (10.68Å/px) were acquired. Tilt series were acquired using Parallel Cryo-Electron Tomography (PACEtomo) in SerialEM using the following settings. Tilt series were acquired at 33000× magnification (2.671 Å/pix) either with an objective lens, a defocus of -3 µm and an objective aperture of 100 µm. The dataset of A/WSN/33 (H1N1) presented in Fig. 1A was acquired in focus with a Volta phase plate and no objective aperture. The tilt series were acquired using a dose symmetric tilt series starting at 0°, with a tilting range from 60° to -60° and increments 3°. Motion correction of acquired movies was done with Motioncor2 (56). Tomograms were reconstructed in IMOD where colloidal gold beads were used as fiducial markers during tilt series alignment (57). Contrast transfer function estimation and correction by phase flipping was performed in IMOD in defocused data. A weighted-back-projection algorithm with a SIRT-like filter 5 was used to reconstruct tomograms after dose-exposure filtering in IMOD and binned three times for figure production.

### Structure modelling and charge calculations

Hemagglutinin models were prepared using AlphaFold3 server (58). Electrostatic charge display was performed in ChimeraX (59). Isoelectric points at different pH were calculated using Prot pi tool, version 2.2.29.152 (https://www.protpi.ch/Calculator/ProteinTool)

### Cell-cell fusion assay

MDCK cells stably expressing HLA-DR_TagRFP_ were cultured in DMEM-GlutaMAX-I medium supplemented with 10% fetal bovine serum (FBS), 1% peniclillin/streptomycin (P/S) and 2.5 µg/ml puromycin at 37°C and 5% CO_2_. Cells were seeded at a density of 1.5 × 10^4^ cells per well in a 24-well plate. Approximately 16 hours post-seeding, cells were transfected with plasmids encoding for GFP and respective IAV HA encoding plasmids H1 (pCDNA3.1_HA_A/WSN/1933_H1N1, UniProt: I4EPC4), H18 (pCDNA3.1_HA_A/flat-faced bat/Peru/03/2010_H18N11). Transfection mixtures were prepared with 50 µl OptiMEM, 1 µg total DNA in a 1:1 ratio of GFP and HA plasmids, and 1.5 µl TransIT-LT1. After incubation of the mixture at room temperature for 30 min, the cell medium was removed and replaced with 200 µl OptiMEM, and the respective transfection mixture was applied dropwise to each well. Cells were incubated at 37°C for five to six hours before the medium was exchanged to complete DMEM. After 24 h, the cell-cell fusion assay was conducted. Transfected cells were washed three times with 500 µl PBS and incubated in 500 µl serum-free DMEM + 2 µg/ml TPCK-Trypsin for 30 min at 37°C to allow for cleavage of HA. Following cleavage, cells were washed once with 500 µl PBS, then incubated in 500 µl complete DMEM adjusted to pH 7.5, 5.5 or 4.5 using citric acid at 37°C for 20 min. After pH treatment, cells were washed once with 500 µl PBS, given 500mL complete DMEM at neutral pH, and incubated at 37°C for 1.5 h to allow for fusion to occur. Cells were then fixed with 250 µl 4% PFA for 15 minutes, washed and stored in 500 µl PBS at 4°C. Before imaging, cells were stained with DAPI at 1:2000 concentration and washed twice with 500 µL PBS. Fluorescence images were acquired by Zeiss CellDiscoverer 7 microscope equipped with an Axiocam 712 camera, using the 5× objective lens. Syncytia were quantified in ImageJ/Fiji using the StarDist plugin to quantify nuclei (60, 61).

### Hemagglutination assay

An aliquot of 1 ml concentrated erythrocytes from a human blood sample was centrifuged at 700 g and 4°C for 7 min, the supernatant removed, and cells resuspended in 500 µl PBS. Centrifugation and PBS washing were repeated a total of three times, and cells were then diluted to a concentration of 1% in PBS. IAV H1N1 WSN and H18N11 were thawed on ice and diluted in PBS to a titer of 1 × 10^7^ ffu/ml. A two-fold serial dilution in PBS was conducted in triplicates for each viral strain on a 96-well plate, and 1 % erythrocytes were added to all wells at a 1:1 volumetric ratio. The plate was incubated at room temperature for 45 min, then examined for hemagglutination and imaged.

## Acknowledgments

We thank the Infectious Disease Imaging Platform (IDIP) at the Center for Integrative Infectious Disease Research Heidelberg and the cryo-EM network at Heidelberg University (HDcryoNET) for support and assistance. The authors gratefully acknowledge the data storage service SDS@hd supported by the Ministry of Science, Research, and the Arts Baden-Württemberg (MWK), the German Research Foundation (DFG) through grant INST 35/1314-1 FUGG and INST 35/1503-1 FUGG.

## Funding

Chica and Heinz Schaller Foundation (Schaller Research Group Leader Programme) (P.C.)

Deutsche Forschungsgemeinschaft (DFG, German Research Foundation): Project number SFB-1129/3 – 240245660 – P19 (P.C.)

DFG Heisenberg program: Project number 537227910 (P.C.)

European Research Council (ERC) (NUMBER 882631—Bat Flu) (M.S)

Excellence Initiative of the German Research Foundation (GSC-4, Spemann Graduate School) (M.S)

Hans A. Krebs Medical Scientist Programme of the Medical Faculty of the University of Freiburg (P.R)

## Author contributions

Conceptualization: SP, PR, PC

Methodology: SP, JR, MKO, RAH, REL, KF, KC, PC

Investigation: SP

Visualization: SP, PC

Funding acquisition: PC, MS, PR

Project administration: PC

Supervision: PC, PR, MS

Writing – original draft: SP

Writing – review & editing: all authors

## Data, code, and materials availability

Cryo-electron tomography data will be deposited in the Electron Microscopy Data Bank (EMDB) and will be available upon publication. Additional data and material related to this publication may be obtained upon request.

