## Supplementary Figures for "MHC-II acts as a fusion-triggering receptor for bat influenza virus"

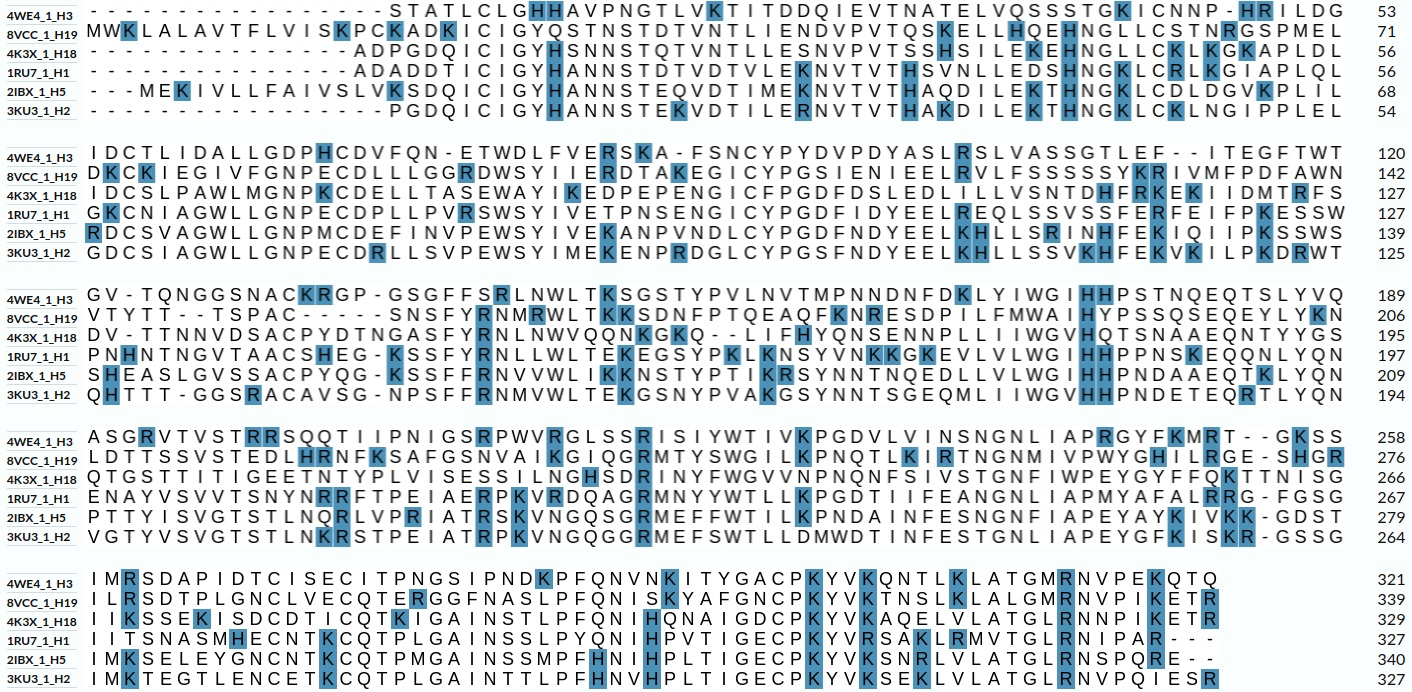

Fig. S1.

**Sequence alignment of different IAV HA subtypes.** Sequence of the HA1 domains from H1, H2, H3, H15, H18, and H19 with indicated PDB codes. Positively charged amino acids are highlighted in blue. Two conserved histidine residues at position 38 and 184 (H3 numbering, red box) were reported to play a role in low pH sensing (25,26).

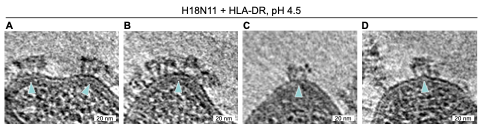

Fig. S2.

**H18N11 HA clusters and membrane curvature.** **A-D.** Slices through cryo-electron tomograms of H18N11 virions incubated with HLA-DR at pH 4.5 showing clusters of HA in postfusion confirmation in regions of high viral membrane curvature (blue arrowheads). Scale bars: 20 nm.

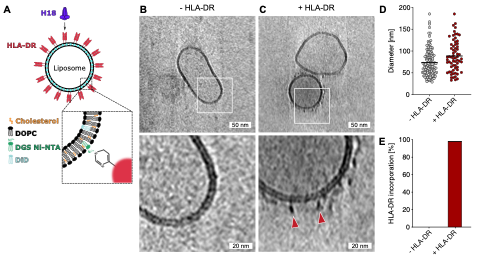

Fig. S3.

**Liposome production.** **A.** Schematric representation of a liposome with bound HLA-DR (red) composed of 52.5% DOPC (black), 40% cholesterol (orange), 5% DGS-Ni-NTA (green) and 2.5% DiD (blue). Bat IAV H18 (purple) uses HLA-DR as entry receptor. **B.** Slice through a cryo-electron tomogram showing a liposome without HLA-DR. Scale bar: 50 nm, scale bar of zoom-in: 20 nm. **C.** Slice through a cryo-electron tomogram showing two liposomes with bound HLA-DR (red arrowheads). Scale bar: 50 nm, scale bar of zoom-in: 20 nm. **D.** Quantification of liposome diameter from cryo-EM overview maps for liposomes without HLA-DR (grey) and with HLA-DR (red). **E.** Percentage of HLA-DR incorporation for liposomes with and without HLA-DR.

**Table S1.**

Liposome composition

| Component | Abbreviation | Molar ratio | Supplier |
| --- | --- | --- | --- |
| 1,2-Di-(9Z-octadecenoyl)-*sn*-glycero-3-phosphocholin | DOPC | 52.5% | Avanti, 850375P |
| Cholesterol | Chol | 40% | Avanti, 700000P |
| 1,2-Di-(9Z-octadecenoyl)-sn-glycero-3-[(N-(5-amino-1-carboxypentyl)iminodiessigsäure)succinyl] | DGS-NTA(Ni) | 5% | Avanti, 790404P |
| 1,1'-Dioctadecyl-3,3,3',3'-Tetramethylindodicarbocyanine | DiD | 2.5% | Biotium, 60014 |
